# Study of Pinelliae Rhizoma hepatotoxicity based on complex network algorithm improvement

**DOI:** 10.1101/2022.11.29.518337

**Authors:** Aijun Zhang, Zhaohang Li, Guanpeng Qi, Ze Xu, Xin Liu, Juman Ma, Zuojing Li

## Abstract

Important protein identification methods based on centrality have reached a high level of accuracy. However, there is a need to improve centrality algorithms because they currently consider the nature of protein–protein interaction (PPI) network topology but not protein properties. To improve the centrality algorithm, we introduce the weighted PageRank algorithm, which represents node importance, and the protein interaction combined_score, which represents PPI network edge importance in the STRING database to construct a weighted PPI network. We constructed yeast protein networks for simulations to validate the improved algorithm. Finally, we studied the hepatotoxicity of Pinelliae Rhizoma by applying the PageRank and Edge Clustering (PEC) algorithm. Our study shows that the PageRank and Edge Clustering algorithm can pre-screen important targets and has superior accuracy, sensitivity, and specificity to other centrality algorithms.

## 1. Background

The concept of network toxicology was first proposed by Changxiao Liu et al. It is a toxicity research method in Chinese medicine developed from network pharmacology. By constructing network models and applying network analysis methods to predict drug toxicity, network toxicology studies the toxic side effects and mechanisms of action on the organism to achieve toxicity reduction and efficacy enhancement^[1]^. The emphasis of the network analysis in network toxicology is predicting the key proteins in the PPI network.

Important proteins are essential for organisms and play important roles in life activities. The identification of important proteins has great value for life sciences and disease treatment. Using computational methods, important proteins at specific locations and under specific conditions can be rapidly predicted^[2]^. Moreover, we can obtain a large amount of protein interaction data from databases. However, the identification of important proteins through current centrality algorithm methods can only reflect the characteristics of nodes, not the importance of a given protein interaction or the protein itself. Therefore, we need an improved algorithm to achieve better prediction^[3]^.

Existing computational-based algorithms for important protein identification in PPI networks have solved some problems. However, the accuracy of important protein identification is still at a moderate level and far from what we would expect^[4]^. Traditional algorithms use one or more parameters and features for analysis, such as degree centrality (DC), betweenness centrality (BC), and closeness centrality (CC), without assigning weights to distinguish the importance of the different parameters. There are also some algorithms that focus simply on the topological features of PPI networks, ignoring the importance of specific proteins in biology and that some false negative or false-positive data may exist in PPI networks. Therefore, we need an algorithm that reflects the importance of nodes, network topology, protein properties and protein interactions^[5]^. The PPI network is a complex and unreliable network with a small-world nature. Since we obtained important protein data with a high percentage of false-positives and false negatives from the database, we should not only combine biological information to eliminate the influence of false-positives and negative influences on the PPI network but also verify the accuracy and reliability of the key protein identification algorithm from different perspectives. Finally, it is necessary to apply the improved algorithm to the study of network toxicology of Chinese medicine.

Pinelliae Rhizoma is a Chinese medicine commonly used in clinical practice. The existing toxicity studies of Pinelliae Rhizoma are mainly focused on four points: mucosal irritation toxicity, hepatorenal toxicity, pregnancy toxicity, and genotoxicity. Currently, studies on its mucosal irritation toxicity are more extensive, and the explanation of this mechanism is more mature^[6]^. However, there are few studies on the toxicity of the other three types of toxicity. Most of these studies are at the stage of observation of general indicators and have not yet reached the molecular level. The understanding of the mechanism of toxicity is not clear and consequently has little significance for the clinical guidance of drug therapy. Most of the studies on Pinelliae Rhizoma combination and detoxification remain focused on its toxic needle crystals^[7]^. Therefore, it is necessary to study the hepatorenal toxicity, pregnancy toxicity and genotoxicity of Pinelliae Rhizoma. We systematically and comprehensively analysed the hepatotoxicity of Pinelliae Rhizoma from the perspective of network toxicology and explored its toxicity mechanism for the purpose of toxicity reduction and efficiency enhancement to better utilize Pinelliae Rhizoma. This study will provide network toxicology support for further studies of the mechanisms of Pinelliae Rhizoma toxicity. The flowchart of technical strategy in present study is illustrated in Fig. 1.

**Fig.1.**
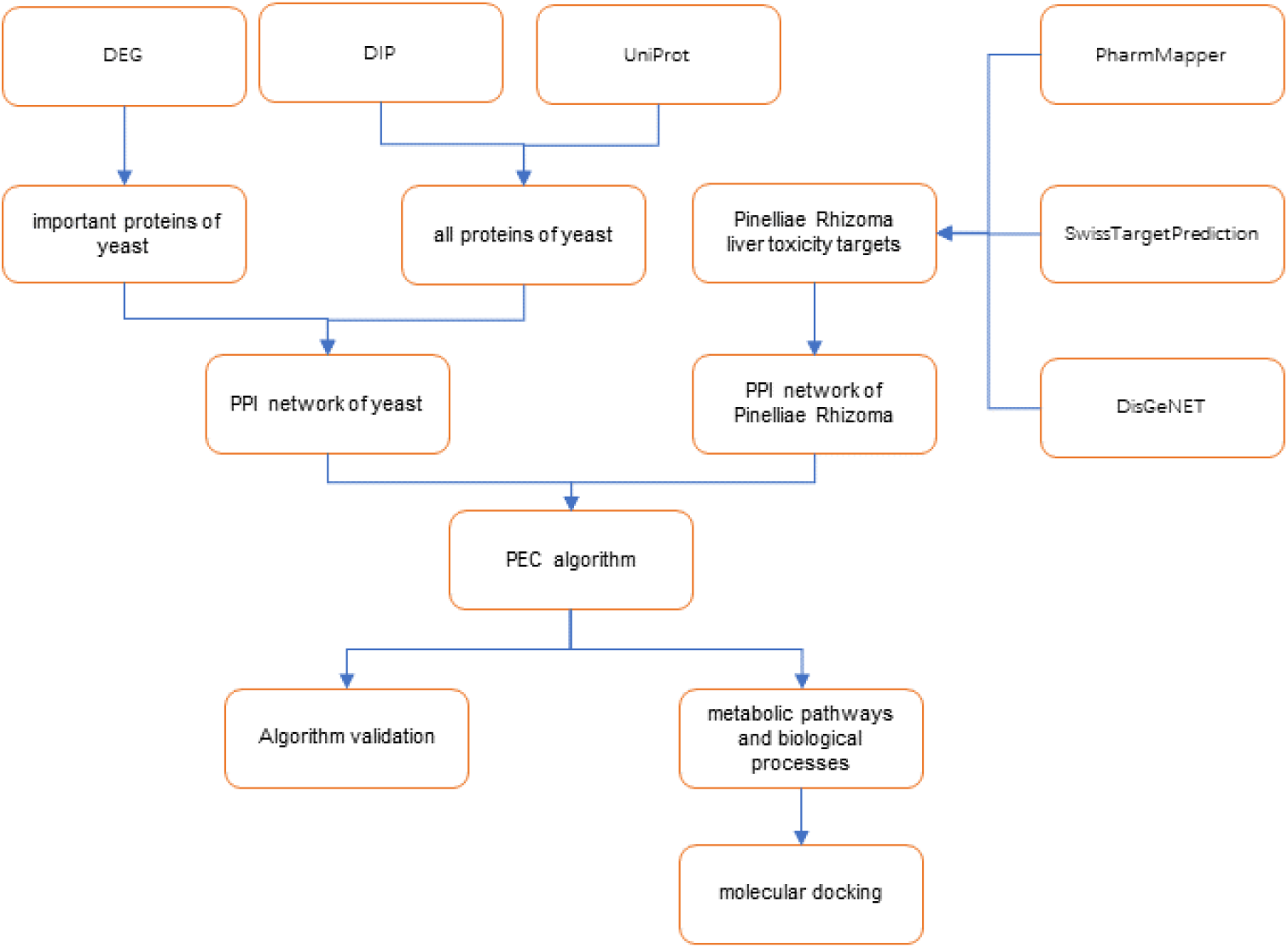
The flowchart of technical strategy in current study

## 2. Method

### 2.1 Algorithm improvement

PageRank is the core algorithm of the Google search engine, which ranks web pages according to their link structure. The importance of a web page on the internet depends on the number and quality of other pages pointing to it. If a page is pointed to by many high-quality pages, then this page is a high-quality page. Its algorithm is as follows:

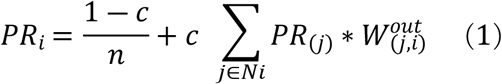

Where n is the total number of nodes. The value of the parameter c depends on the specific situation. Initially, assign an initial PageRank value *PR_i_*(0) to all objects, indicating the probability that the object is connected, and meet 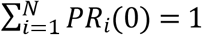. Then iteratively assign the PR value of each object to the object it points to until all PR values converge. The last PR value indicates the importance of the object. 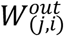 is the link weight ratio from j to i, 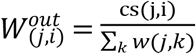, cs(j,i) is the scoring value of the relationship between the corresponding node j and node i in the string database, ∑*_k_ w*(*j,k*) is the sum of link weights of all out-degree of node j ^[8]^.

Traditional centrality algorithms describe only edge or point properties to predict important targets without first integrating the importance of individual edges and points^[9]^. In contrast, our method does integrate the importance of both edges and points. First, we represent the importance of an edge uv by the number of triangles it forms in the network. We consider that the more triangles an edge forms in the network, the more important that edge is. Second, the importance of nodes represented by each protein is different in PPI networks. Therefore, we introduce the weighted PageRank algorithm to assign a corresponding weight to each node. The formula to calculate the importance of edge uv is as follows:

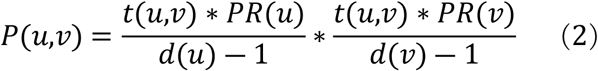

where t(u,v) denotes the number of edges uv forming triangles in the network. PR(u) denotes the PageRank value of point u. d(u) denotes the degree of point u.

In the PPI network, not only is the importance of each protein different, but the protein–protein interaction relationships are also different. Therefore, we used the protein interaction score combined_score from the STRING database to indicate the strength of each protein–protein interaction relationship to construct a weighted PPI network. The PEC algorithm is as follows:

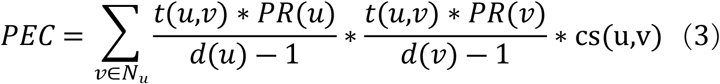

where denotes the number of edges t(u, v) uv that form a triangle in the network. PR (u)denotes the PageRank of point u. d(u) denotes the degree of point u. N_u_ denotes the set of edges that already exist in the network. cs(u,v) is the scoring value of the strength of the relationship combined_score, between the corresponding nodes u and v, in the STRING database.

### 2.2 Algorithm validation

The protein interaction data and important protein data of the yeast *Saccharomyces cerevisiae* are the most complete and reliable of all species to date. Karagoz et al.^[10]^ built a highly reliable and almost complete yeast PPI network from several noncommercial databases. Estrada et al.^[11]^ applied subgraph centrality and five other classical centrality algorithms to predict key targets in yeast PPI networks. In our experiment, we build an important proteins dataset and an all-proteins dataset for yeast by finding and extracting yeast protein data from multiple databases. Then, we build a yeast PPI network from the yeast all-proteins dataset and apply the PEC algorithm to the prediction of important proteins in the yeast PPI network. Finally, the accuracy of PEC algorithm identification and other indices were obtained by comparison with an established important yeast protein dataset.

The important proteins of yeast were found in the Database of Essential Genes (DEG)^[12]^. Then, the Database of Interacting Proteins (DIP)^[13]^ and UniProt database were searched for yeast proteins, and their union was taken to obtain all the proteins of yeast. Then, the UniProtKB data corresponding to the proteins were transformed into the corresponding gene names. Finally, the PPI network of yeast was obtained by importing all of the yeast genes into the STRING database^[14]^.

### 2.3 Hepatotoxicity study of Pinelliae Rhizoma

The ingredients of Pinelliae Rhizoma were identified in the Traditional Chinese Medicine Systems Pharmacology Database and Analysis Platform (TCMSP) (https://tcmsp-e.com/)^[15]^ and then screened for ingredients with oral bioavailability (OB) > 30% and drug-like (DL) score > 0.18^[16]^. The TCMSP, CTD^[17]^ and ADMETlab 2.0 databases^[18]^ were searched to determine the hepatotoxicity of these components, and those with hepatotoxicity were screened. The hepatotoxic components of Pinelliae Rhizoma were then imported into the PharmMapper database^[19]^ and SwissTargetPrediction database for target prediction, and all targets obtained from both databases were combined and deduplicated. The DisGeNET database collects a large number of variants and genes associated with human diseases, and hepatotoxicity targets were obtained from DisGeNET. The relevant targets were intersected with the targets obtained from the above two databases and used as potential hepatotoxicity targets of Pinelliae Rhizoma. The potential hepatotoxicity targets of Pinelliae Rhizoma were imported into the STRING database to obtain the PPI network.

The PPI network was calculated by the PEC algorithm, and the targets with higher scores were the more important targets in the PPI network. To further investigate the synergistic effects of Pinelliae Rhizoma hepatotoxicity target clusters, Metascape^[20]^ was used for Gene Ontology (GO) enrichment analysis, Kyoto Encyclopedia of Genes and Genomes (KEGG) pathway analysis and WikiPathways analysis. Finally, the accuracy of the toxicity target prediction was verified by using iGEMDOCK2.1 software to perform molecular docking.

## 3. Results

### 3.1 Algorithm validation results

First, the yeast PPI network was constructed: 1110 important proteins of yeast were found in the DEG database. A total of 5,058 proteins were obtained by searching for yeast proteins in the DIP database and UniProt database and taking their intersection. Then, the yeast protein UniProtKB data were converted into corresponding gene names, and all the genes were imported into a STRING database multiple times, merged and deduplicated to obtain the complete PPI network of yeast. The yeast PPI network contains 22798 protein–protein interaction edges and 5058 yeast proteins, including 1110 important proteins.

The PEC algorithm was programmed in the R programming language, and the yeast PPI network was calculated by using R to predict the key targets. The PEC algorithm was evaluated by comparison with three algorithms: closeness centrality (CC), betweenness centrality (BC), and degree centrality (DC). Since there were 1110 important proteins in the PPI network, the top 1%, 2%, 5%, 10%, 15%, 20%, and 25% of proteins were taken as the candidate sets of important proteins. By intersecting these sets with the yeast important protein data, the true important proteins among the candidate important proteins can be derived. As shown in Fig. 2, the number of important proteins correctly predicted by the PEC algorithm is significantly improved compared with that predicted by other algorithms.

**Fig.2.**
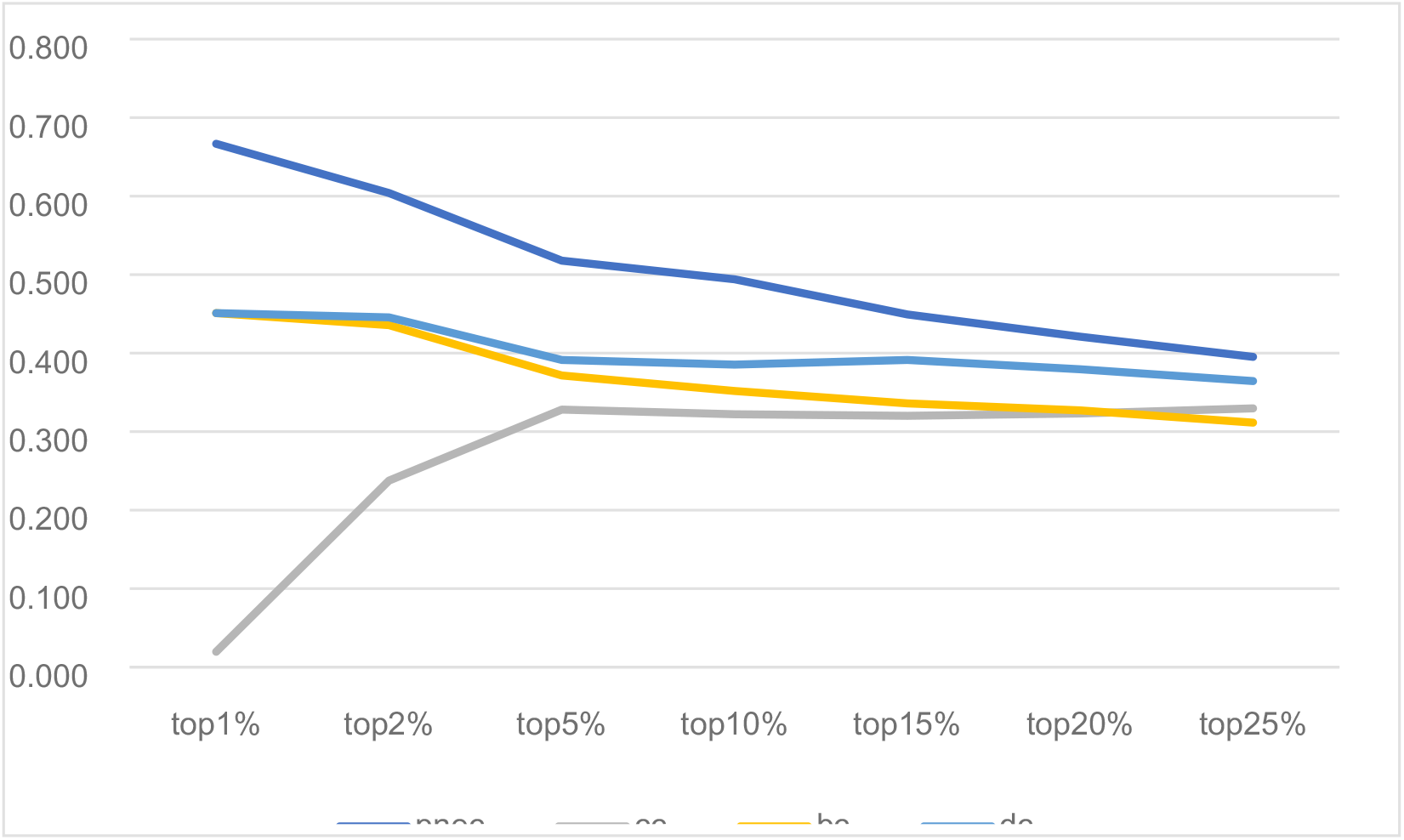
Accuracy of each algorithm in predicting important proteins at different levels

To evaluate the performance of the new algorithm PEC in predicting important proteins, several commonly used indicators in medical tests are introduced. To understand the concept of each indicator, four commonly used statistical measures are given, and their respective meanings are explained in Table 1.

**Table 1.** four commonly used statistical terms

| Measures | Explain or Formula |
| --- | --- |
| true positives ( TP ) | Important proteins are correctly identified as Important proteins |
| false positives ( FP ) | non-Important proteins are incorrectly identified as Important proteins |
| true negatives ( TN ) | non-Important proteins are correctly identified as non-Important proteins |
| false negatives ( FN ) | Important proteins are incorrectly identified as non-Important proteins |
| sensitivity (SN) | $SN = \frac{TP}{TP + FN}$ |
| specificity (SP) | $SP = \frac{TN}{TN + FP}$ |
| positive predictive value (PPV) | $PPV = \frac{TP}{TP + FP}$ |
| negative predictive value (NPV) | $NPV = \frac{TN}{TN + FN}$ |
| F-measure (F) | $F = \frac{2 \times SN \times PPV}{SN + PPV}$ |
| accuracy (ACC) | $ACC = \frac{TP + TN}{P + N}$ |

On this basis, six statistical metrics commonly used in medicine, sensitivity (SN), specificity (SP), positive predictive value (PPV), negative predictive value (PPV), F-measure (F) and accuracy (ACC), were calculated^[9]^. The results are shown in Table 2, indicating that the PEC algorithm has improved in various measures.

**Table 2.** Comparison of evaluation index values for each algorithm

|  | **SN** | **SP** | **PPV** | **NPV** | **F** | **AAC** |
| --- | --- | --- | --- | --- | --- | --- |
| **PEC** | 0.422 | 0.856 | 0.422 | 0.856 | 0.422 | 0.769 |
| **DC** | 0.381 | 0.843 | 0.381 | 0.843 | 0.381 | 0.749 |
| **BC** | 0.329 | 0.829 | 0.329 | 0.829 | 0.329 | 0.728 |
| **CC** | 0.329 | 0.829 | 0.329 | 0.829 | 0.329 | 0.728 |

Based on false positive and true positive, false positive rate and true positive rate were used to draw ROC curve as the ordinate and abscissa. The ROC curve of the subject’s working characteristics is often used to evaluate the quality of the binary classification system, which is the most objective and effective method so far. Each point on the ROC curve reflects the receptivity to the same signal stimulus. AUC, the area under the ROC curve, is often used as an important indicator to analyse the recognition rate. The larger the AUC, the better the algorithms performance. As shown in Figure 3, the ROC curves were compared with PEC, CC, BC and DC. and that the performance of PEC algorithm were founded better than the other three algorithms.

**Fig.3.**
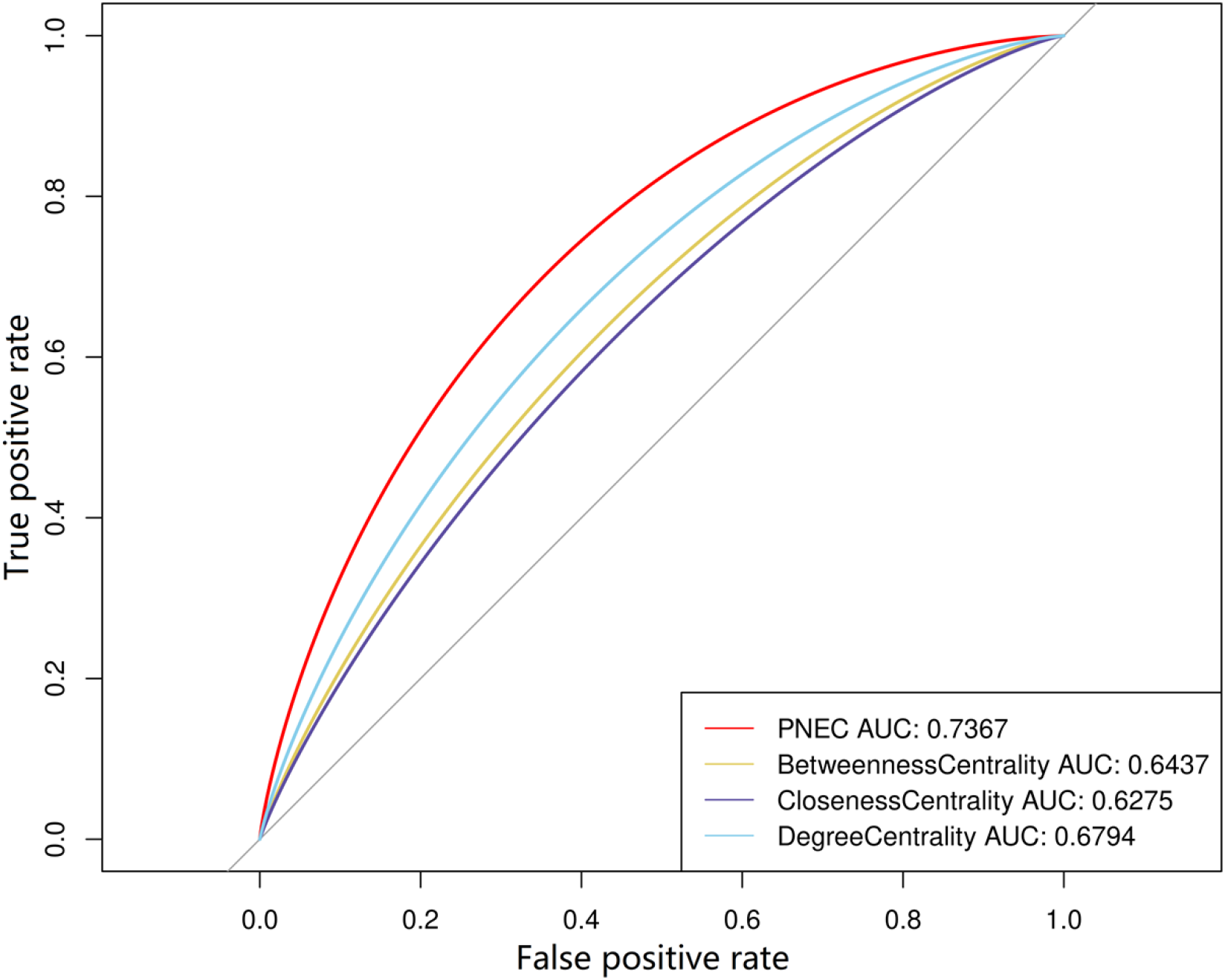
ROC curve and AUC value

In order to analyse the PEC algorithm more carefully, the Jackknife curve were introduced to analyse advantages of it. The abscissa represents the predicted number of key proteins, and the ordinate represents the actual number of proteins in the predicted number of key proteins. The larger the area under the curve, the higher the recognition rate of key proteins. As shown in Figure 4, the recognition rate of key proteins of PEC is higher than that of the other three.

**Fig. 4.**
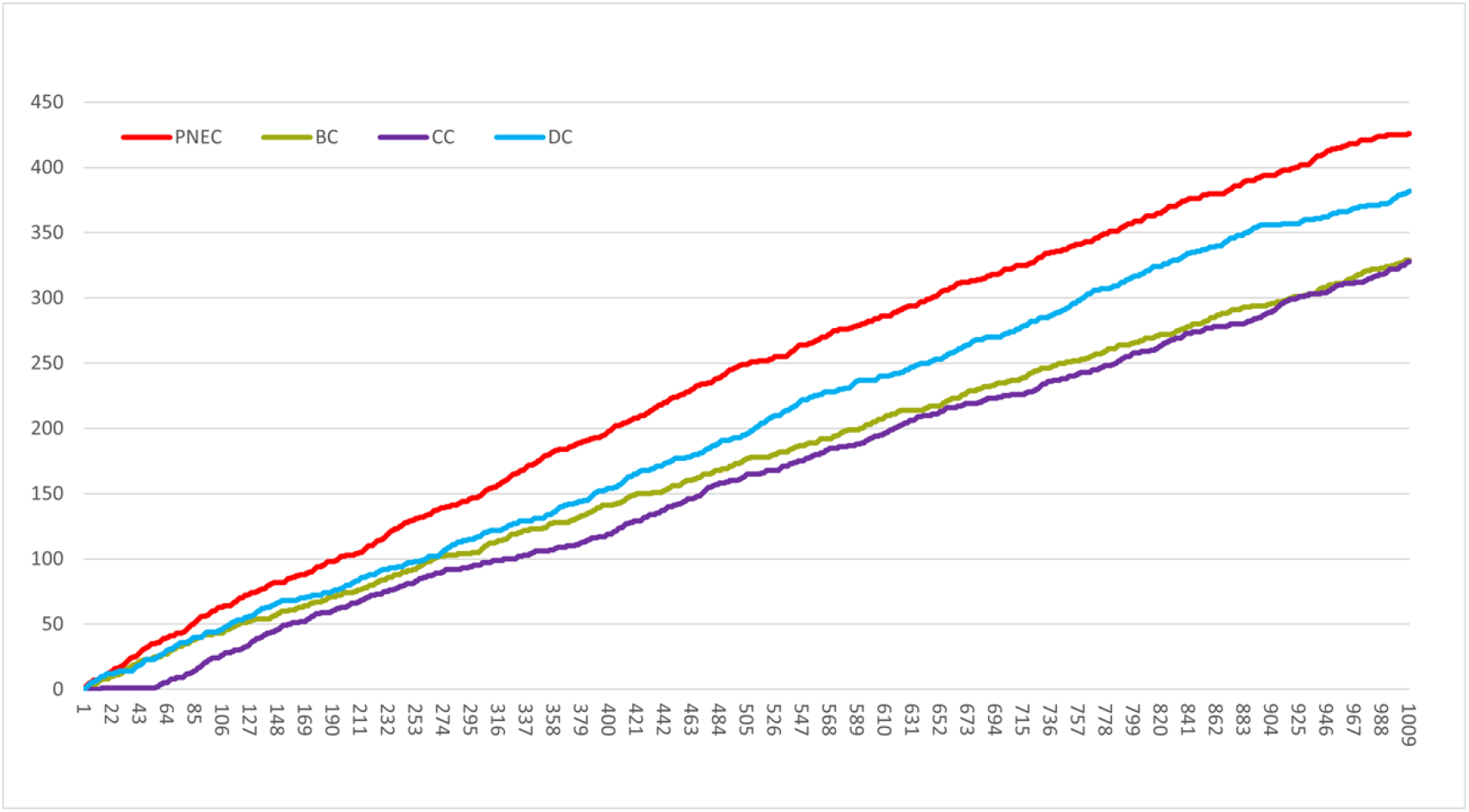
the Jackknife curve

### 3.2 Results of the Pinelliae Rhizoma liver toxicity component collection

A total of 116 ingredients of Pinelliae Rhizoma were found from the TCMSP, and those with OB > 30% and DL > 0.18 were selected for determination of their hepatotoxicity in the CTD database, ADMETlab database and TCMSP database. Finally, 13 ingredients with hepatotoxicity were screened, as shown in Table 3.

**Table 3.**
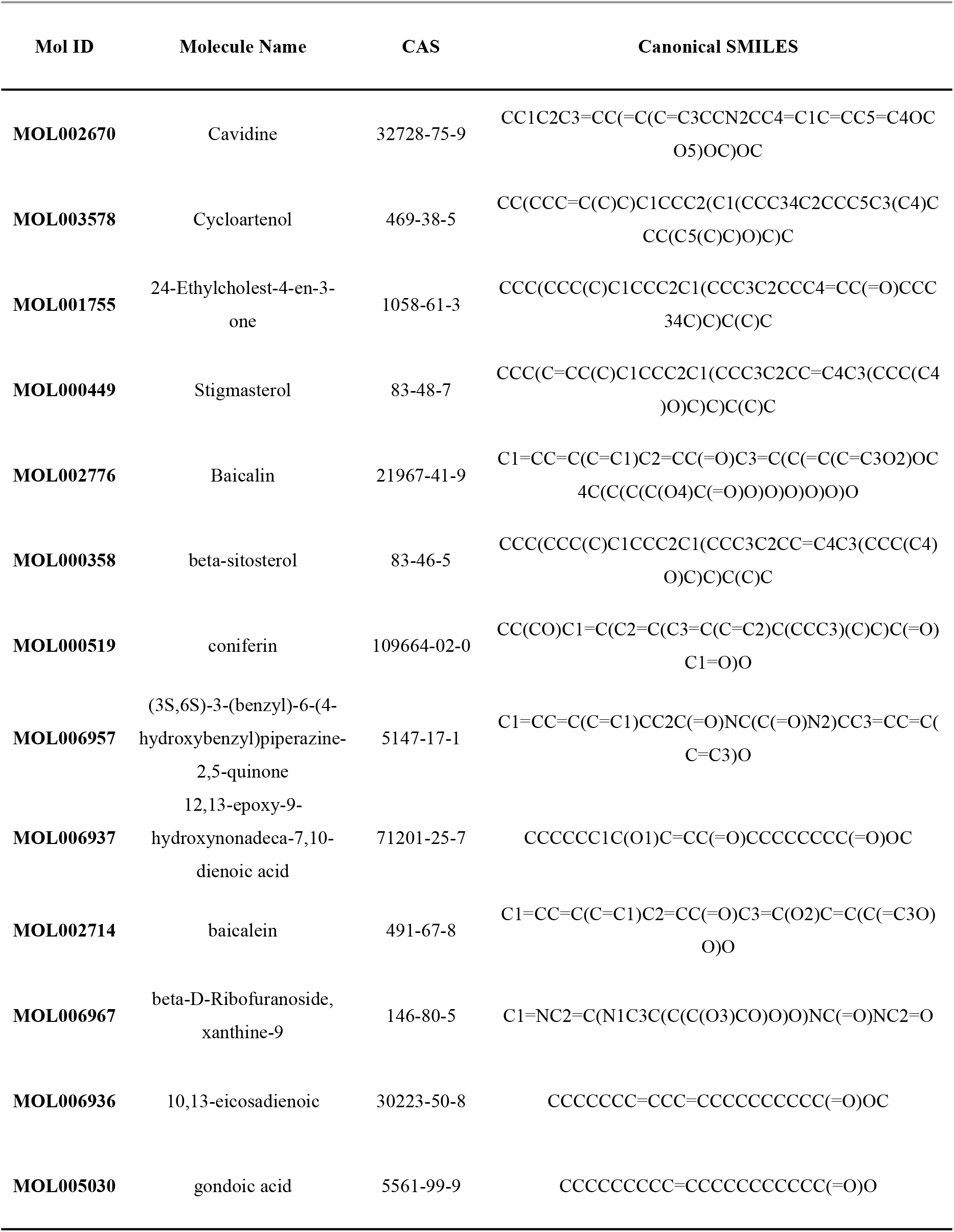
Hepatotoxic components of Pinelliae Rhizoma

To predict the targets of these Pinelliae Rhizoma active ingredients, the 13 ingredients were entered into two databases, PharmMapper and SwissTargetPrediction, for target prediction. A total of 430 targets were found in the PharmMapper database, and 346 targets were found in the SwissTargetPrediction database. The targets obtained from both databases were combined and deweighted to obtain a final set of 660 targets for Pinelliae Rhizoma. A total of 1019 liver disease-related targets were found in the DisGeNET database and intersected with the 660 Pinelliae Rhizoma targets to obtain 153 Pinelliae Rhizoma liver toxicity targets. The 153 liver toxicity targets were imported into the STRING database to obtain a PPI network, which contains 153 nodes and 1889 interaction edges.

### 3.3 Results of the Pinelliae Rhizoma hepatotoxicity study

The PPI network was imported into R, and the key targets were predicted using the PEC algorithm. It was concluded that STAT3, PTPN11, and other targets ranked highly and were the key targets of the PPI network. The top 10% of the key targets are shown in Table 4.

**Table 4.** Top 10% of key targets

| Gene name | UniProt ID | Protein names |
| --- | --- | --- |
| STAT3 | P40763 | Signal transducer and activator of transcription 3 |
| PTN11 | Q06124 | Tyrosine-protein phosphatase nonreceptor type 11 |
| PGH2 | P35354 | Prostaglandin G/H synthase 2 |
| MMP9 | P14780 | Matrix metalloproteinase-9 |
| MMP3 | P08254 | Stromelysin-1 |
| MMP2 | P08253 | 72 kDa type IV collagenase |
| MMP1 | P03956 | Interstitial collagenase |
| MK08 | P45983 | Mitogen-activated protein kinase 8 |
| MK03 | P27361 | Mitogen-activated protein kinase 3 |
| MK14 | Q16539 | Mitogen-activated protein kinase 14 |
| MK01 | P28482 | Mitogen-activated protein kinase 1 |
| VGFR2 | P35968 | Vascular endothelial growth factor receptor 2 |
| EGFR | P00533 | Epidermal growth factor receptor |
| CASP3 | P42574 | Caspase-3 |
| B2CL1 | Q07817 | Bcl-2-like protein 1 |
| AKT1 | P31749 | RAC-alpha serine/threonine-protein kinase |

The Metascape platform was used to analyse 153 potential toxicity targets of Pinelliae Rhizoma for metabolic pathways and biological processes. KEGG signalling pathways, GO Biological Processes, and WikiPathways with p value< 0.01, a minimum gene number of 3 and an enrichment factor > 1.5 were selected for analysis. 47 important metabolic pathways and biological processes were found. Combining the data of ingredients and proteins, as shown in Fig.5, the ingredient-target-pathway network of Pinellia ternata hepatotoxicity was constructed.

**Fig. 5.**
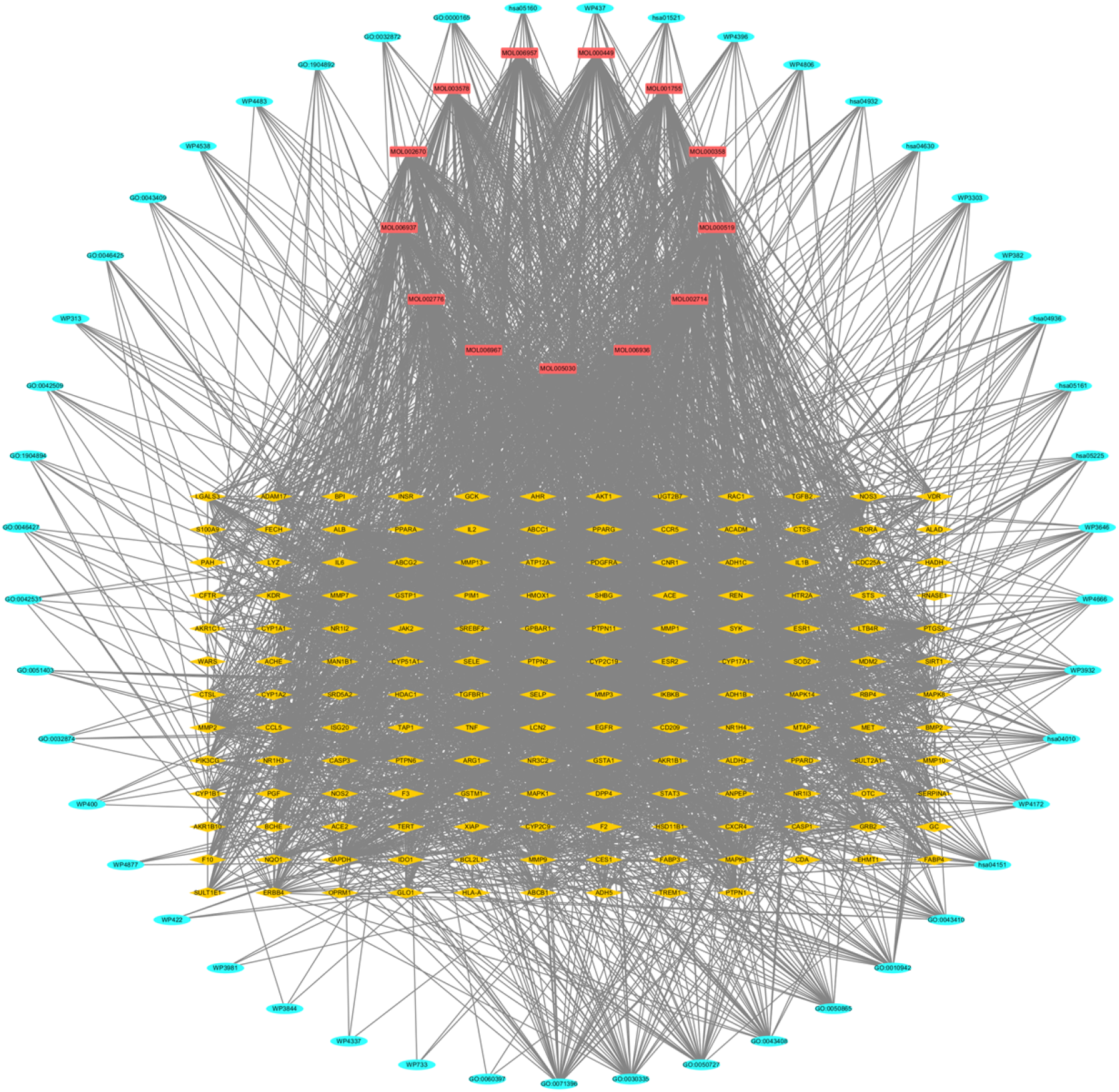
ingredient-target-pathway network of Pinelliae Rhizoma

The metabolic pathways and biological processes, as shown in Fig.6, mainly included MAPK cascade regulation, the JAK-STAT regulation of receptor signalling pathway, the RAC1/PAK1/p38/MMP2 pathway, the PI3K-Akt signalling pathway, and the EGF/EGFR signalling pathway.

**Fig. 6.**
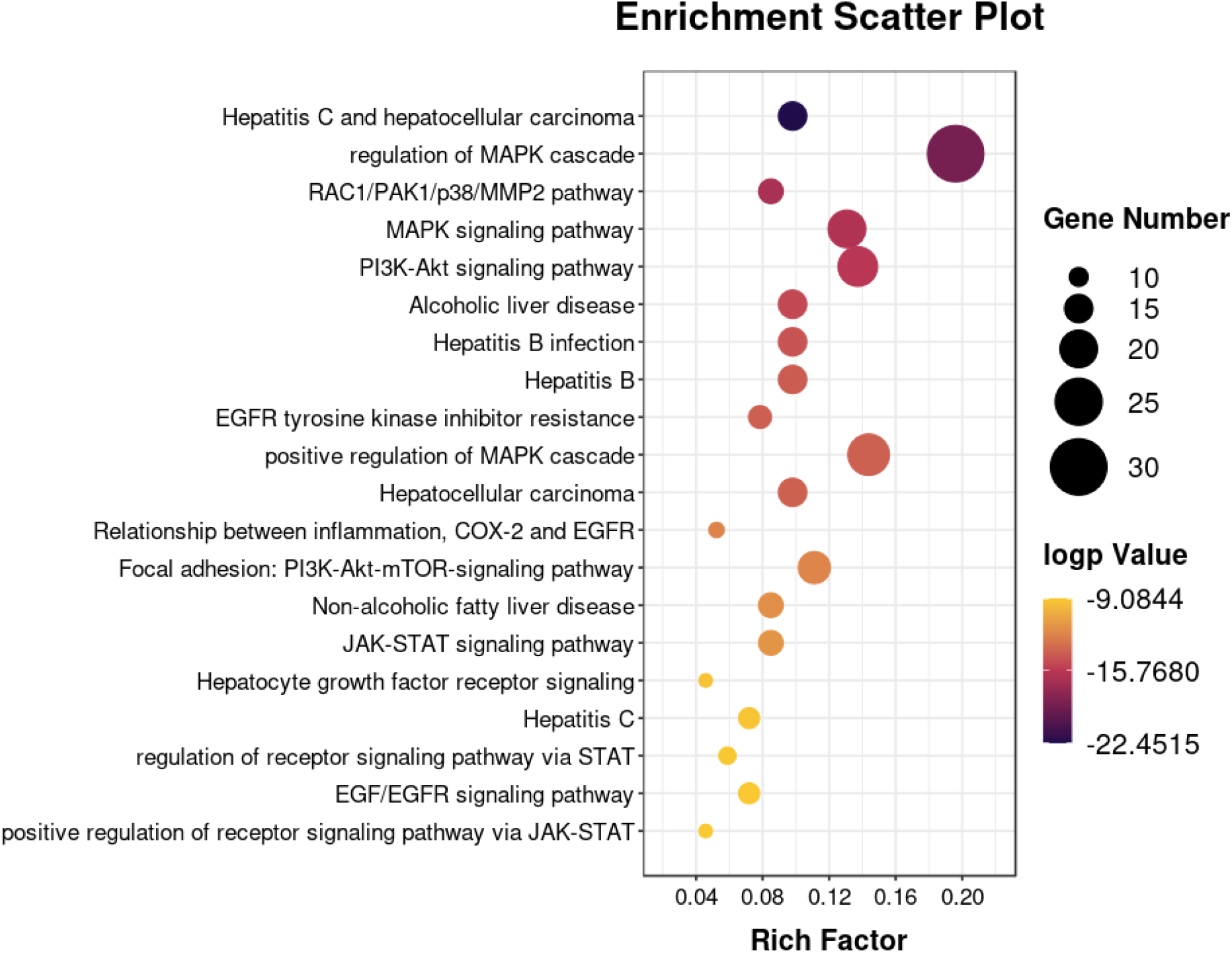
Metabolic pathway and biological process enrichment analysis

MAPK1/3 plays a role in autophagy and is implicated in lipid metabolism. MAPK 1/3 affects hepatic lipid metabolism by stimulating ATG7-dependent autophagy^[21]^. Several studies have also shown that mir-224 plays a role in the proliferation, migration, invasion and apoptosis resistance of hepatocellular carcinoma cells through direct binding to its target genes, and the MAPK 1 gene has been shown to be an important target of mir-224^[22]^.

STAT3 is a key regulatory gene in the JAK/STAT signalling pathway, and activation of the JAK/STAT signalling pathway increases the level of p-STAT3 protein in cells, which is essential for rapidly regulating gene transcription levels and influencing the proliferative and invasive activity of tumour cells. STAT3 is aberrantly highly expressed in all major human malignancies and mainly regulates downstream genes, thereby regulating cell proliferation, apoptosis, and malignant transformation. STAT3 expression in signalling pathways is closely related to hepatocellular carcinogenesis^[23]^.

MMP-2 and MMP-9 are proteolytic enzymes that degrade type IV collagen, and they can disrupt the balance of extracellular matrix degradation, thus allowing cancer cells to break through the extracellular matrix and basement membrane and achieve local invasion and metastasis. MMP-2 can activate MMP-9, which then activates other matrix metalloproteinases to promote extracellular matrix degradation^[24]^. The expression of MMP-2 was significantly increased in tumour tissues with portal vein infiltration, intrahepatic metastasis or postoperative recurrence, suggesting that MMP-2 plays an important role in the infiltration and metastasis of human hepatocellular carcinoma^[25]^. The expression of MMP-9 was significantly higher in hepatocellular carcinoma with invaded peritoneum than in hepatocellular carcinoma with uninvaded peritoneum, and its mRNA was highly expressed in tissues at the tumour infiltration boundary. Studies have shown that MMP-9 is closely associated with invasion and metastasis in hepatocellular carcinoma^[26]^.

AKT1 is an upstream protein of the PI3K/Akt signalling pathway, and its phosphorylation process activates the pathway and regulates tumour cell proliferation and apoptosis^[27]^. Studies have shown that the PI3K/Akt signalling pathway promotes tumour cell proliferation and accelerates tumour cell infiltration and migration. AKT1 is highly expressed in hepatocellular carcinoma tissues and plays a key role in the development and progression of hepatocellular carcinoma. In addition, AKT1 plays a unique role in the development of inflammation, cell proliferation, migration and fibrosis during alcoholic liver injury. AKT1 is also involved in ethanol- and LPS-mediated liver fibrosis^[28]^.

EGFR (epidermal growth factor receptor) is a transmembrane receptor tyrosine kinase that can be activated by a variety of ligands, activating multiple signalling pathways that primarily control proliferation, differentiation, and survival. The EGFR signalling axis has been shown to play a key role in liver regeneration, cirrhosis and hepatocellular carcinoma following acute and chronic liver injury, highlighting the importance of EGFR in the development of liver disease. A recent study showed that EGFR is upregulated in hepatic macrophages and plays a pro-oncogenic role in human hepatocellular carcinoma and mouse hepatocellular carcinoma models. Thus, the role of EGFR in liver disease appears to be more complex than expected^[29]^.

### 3.4 Analysis of molecular docking results

Molecular docking is the process of placing ligand molecules at receptor active sites, simulating the interaction of ligands and receptors through computer simulation experiments, and finding their optimal binding mode. Using iGemdock2.1 molecular docking software, we selected the top 16 Pinelliae Rhizoma targets as receptors and molecularly docked these targets to the Pinelliae Rhizoma liver toxicity components that are predicted to act on them. The lower the docking energy value is, the stronger the interaction between the ligand and receptor. In iGEMDOCK 2.1 software, a docking energy value less than −90 is considered a strong interaction. Among the 150 docking results, 71.33% were considered strong interactions, as shown in Fig. 5, which suggests that the toxic components of Pinelliae Rhizoma have strong effects on these 16 targets.

**Fig. 5.**
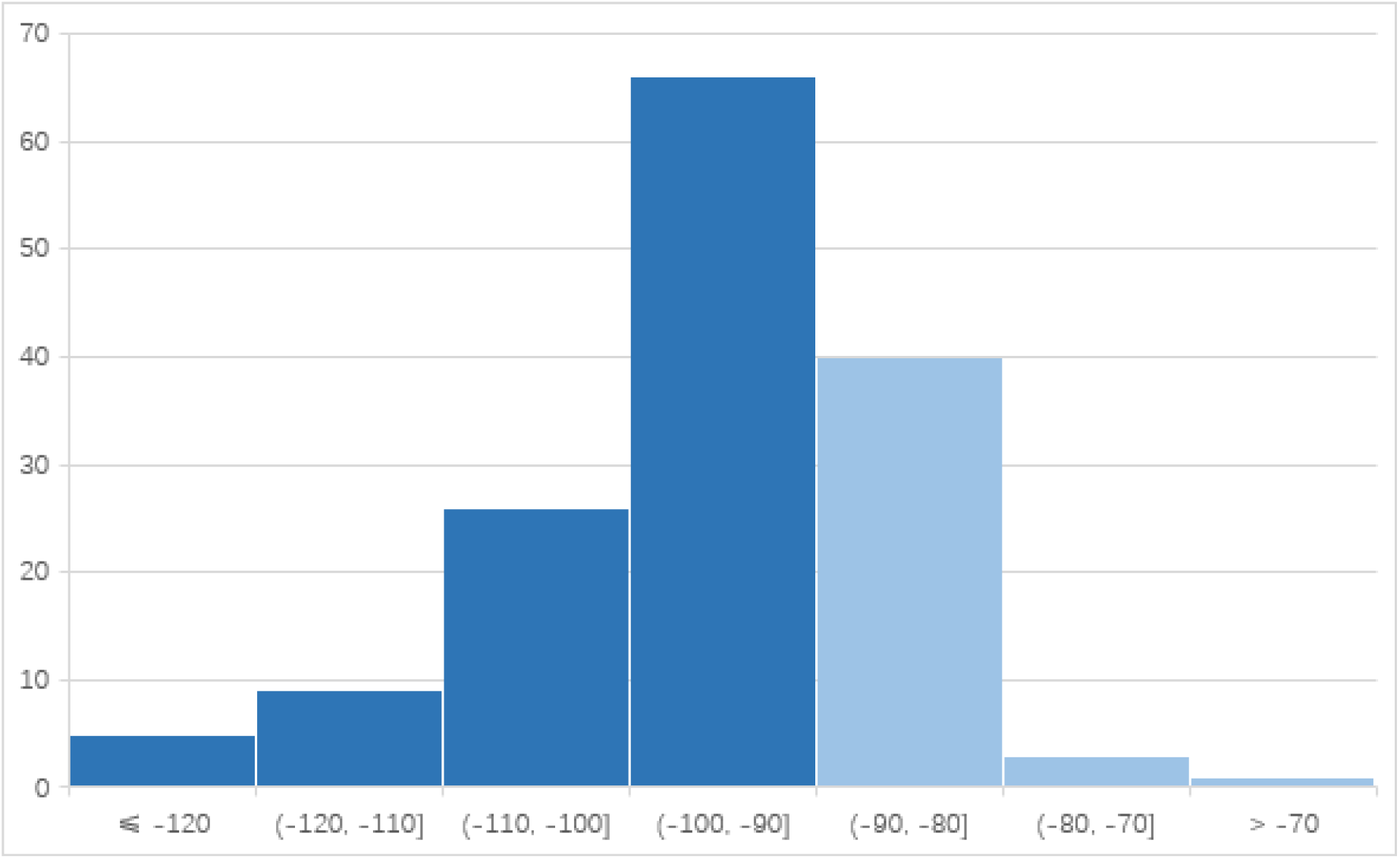
iGEMDOCK molecular docking score statistics

## 4. Conclusion

Network toxicology is a novel concept in the study of the toxicity of herbal medicines. Using the network pharmacology approach to study the toxicity of herbal medicines, the data from databases can be extracted in large quantities, and the datasets can be manipulated directly in the network. The challenges faced by network toxicology and network pharmacology are similar. For example, the database information is limited as it cannot accurately reflect the status of targets and contains some false negative and falsepositive data, and the existing algorithms have a low accuracy of key target prediction.

The following aspects are investigated in this paper. First, we conducted research related to complex networks, improved the algorithm based on the original important protein identification algorithm, introduced the PageRank centrality algorithm and the STRING database protein interaction relationship combined_score, proposed the new algorithm PEC, implemented the new algorithm in R language, and verified the accuracy of the algorithm prediction using brewer’s yeast protein networks. Compared with other centrality algorithms, the PEC algorithm has higher prediction accuracy and is superior in sensitivity, specificity and six other aspects. Second, we conducted a toxicological study of Pinelliae Rhizoma using a network toxicology approach to screen Pinelliae Rhizoma components and find potential toxic targets, and we analysed the toxicological mechanism of action of Pinelliae Rhizoma through protein interaction network construction, pathway analysis, and molecular docking analyses. Hepatotoxicity-related experimental studies of Pinelliae Rhizoma are rare, and one study on the hepatotoxicity of Pinelliae Rhizoma aqueous extract fractions in mice^[30]^ found that higher doses of the Pinelliae Rhizoma aqueous extract fraction resulted in varying degrees of oedema, steatosis, and partial punctate necrosis of hepatocytes.

In this paper, we found that the toxic effects of Pinelliae Rhizoma on the liver are produced through multiple components, targets and pathways, mainly affecting liver lipid metabolism and playing an important role in the proliferation, migration and invasion of liver cancer cells.

## Acknowledgments

This work was supported in part by the National Natural Science Foundation of China (No.41310164).

